# Responses to mechanically and visually cued water waves in the nervous system of the medicinal leech

**DOI:** 10.1101/137588

**Authors:** Andrew M. Lehmkuhl, Arunkumar Muthusamy, Daniel A. Wagenaar

## Abstract

**Summary statement:** Cues from water movement help aquatic predators find their prey. We study how the nervous system of the medicinal leech processes visual and mechanical information derived from surface waves.

**Abstract:** Sensitivity to water waves is a key modality by which aquatic predators can detect and localize their prey. For one such predator, the medicinal leech, *Hirudo verbana*, behavioral responses to visual and mechanical cues from water waves are well documented. Here, we quantitatively characterized the response patterns of a multisensory interneuron, the S cell, to mechanically and visually cued water waves. The frequency dependence of the S-cell response matched the behavioral response well, in that sensitivity was higher for low frequencies in the visual modality and for high frequencies in the mechanical modality. We demonstrated that neither the cephalic ganglia nor the tail brain is required for the S cell to respond to visually cued water waves. The direction of spike propagation within the S- cell system did follow the direction of wave propagation under certain circumstances, but it is unlikely that downstream neuronal targets can use this information. In terms of overall firing rate, the S cell response was not direction selective. Accordingly we propose a role for the S cell in the detection of waves but not in the localization of their source.

## Introduction

Aquatic predators utilize a wide variety of sensory modalities to localize their prey. Some use modalities that are also available to predators in other environments, such as the larvae of the gray treefrog, *Hyla versicolor*, which use chemosensation (Troyer and Turner, 2015), and toothed whales, which use echolocation (Wilson et al., 2007). Others rely on modalities that are only available under water, such as electric eels, which use electroreception (Hagiwara et al., 1965). Yet others rely on the detection of physical disturbances to the water created by their prey. Such indirect detection of prey is widespread through the animal kingdom and has been described in arrow worms (Horridge and Boulton, 1967), insects (Lang, 1980), spiders (Bleckmann and Barth, 1984), toads (Elepfandt, 1984), dogfish (Russell and Roberts, 1974), seals (Dehnhardt et al., 1998; Zimmer, 2001). The medicinal leech, *Hirudo verbana*, is also in this category and utilizes water surface waves to guide predation (Young et al., 1981).

Water waves provide multiple forms of sensory information that can be used to interpret the nature of those waves and—more importantly—the agent responsible for causing them. When a prey animal moves through the water, pressure waves emanate from its location and provide directional cues to predators. Prey animals may also generate surface waves that propagate centrifugally and cause detectable water movement at a distance. Propagating surface waves also cause the refraction of sunlight by the water surface to fluctuate. The resulting alternation of light and shadow cast into the water yields visual cues that can be used as a directional cue.

Medicinal leeches can sense water motion using mechanosensory hairs distributed in a grid along their bodies. These “Sensillar Movement Receptors (SMRs)” provide directional information about the source of surface waves (Young et al., 1981). Additionally, leeches can use visual information (Carlton and McVean, 1993; Dickinson and Lent, 1984) using an array of visual sensilla colocalized with the mechanosensory hairs and several pairs of simple eyes located in the head (Kretz et al., 1976; Röhlich and Török, 1964). The visual sensilla, of which there are 14 pairs in each of the animal’s 21 midbody segments, appear especially well-positioned to detect the direction of passing light and shadows, but leeches do not have image-forming eyes that could be used to see prey directly.

When both mechanical and visual cues are available, adult leeches preferentially use mechanical cues (Harley et al., 2011), but in the absence of mechanical cues, they can still localize their prey using visual cues alone. Although several neurons have been identified as downstream targets of the sensillar photoreceptors (Kretz et al., 1976), the neuronal circuits that allow leeches to discriminate between different visual or mechanical cues and to assess the direction of the surface waves that give rise to those cues remain unknown.

Initiation of hunting behavior requires the combination of sensory information from multiple segments to form the basis of a single ultimate decision. One neuron that may be hypothesized to play a role in this process is the S cell, which occurs as a single unpaired neuron in each of the leech’s 21 segmental ganglia (Laverack, 1969). Each S cell sends axons into the anterior and posterior medial connectives, where they form electrical synapses with the S cells of neighboring ganglia (Frank et al., 1975). The coupling between adjacent S cells is so strong that all the S cells together form a single electric syncytium (hence the name “S” cell). The S-cell system has the fastest conduction velocity of all interganglionic connections in the leech (Frank et al., 1975; Gardner-Medwin et al., 1973; Laverack, 1969). Accordingly, it was long believed that the S-cell system played a central role in escape responses. However, Sahley et al. (1994) demonstrated that the S-cell system is neither necessary nor sufficient for the initiation of whole-body shortening, a key escape response of the leech, although it is activated during sensory-induced shortening and is required for normal plasticity of the shortening response. The S cell is known to respond to sensory stimuli of multiple modalities, including both visual and mechanical modalities (Bagnoli et al., 1973).

Here, we study the S cell’s response to mechanical and visual stimuli associated with water waves. We find that the S cell’s responses to stimuli of different frequency closely matches the previously reported success rate of hunting behavior (Harley et al., 2011), but that the S cell does not encode directional information about approaching waves.

## Methods

### Animal care

Adult Medicinal leeches (*Hirudo verbana*) were obtained from Niagara Medicinal Leeches (Westbury, NY, USA) and maintained according to methods described by Harley et al. (2011). Briefly, leeches were maintained in tanks of 20–40 individuals in a temperature controlled room held at 16°C with a 12h:12h light:dark cycle. All leeches used in this study had fasted for 2–4 months and weighed 1.0–2.7 g at the time of experiments.

### Electrophysiology

Leeches were anesthetized in ice-cold saline (Muller et al., 1981) and immobilized on slabs of transparent DPMS (Sylgard 184, Dow Corning) to allow access to the ventral ganglia. For experiments involving visual cues, leeches were pinned dorsal side down to allow visualization and electrode access from above and stimulation from below. The body wall was opened up along the ventral midline to expose the nerve cord between ganglia 8 and 12. For experiments involving mechanical cues, leeches were pinned dorsal side facing upward and the body wall was opened up along the dorsal midline. For both visual and mechanical experiments, the lateral roots of ganglia 9 through 11 were cut and those three ganglia were immobilized using fine tungsten pins on a thin, rectangular slab of PDMS. Action potentials in the S cell were recorded extracellularly using two *en-passant* suction electrodes applied to the ventral nerve cord on either side of ganglion 10. (In some experiments, we recorded anteriorly and posteriorly to a different ganglion, in which case the lateral roots of that ganglion and its immediate neighbors were cut rather than those of ganglia 9– 11.) Signals were amplified using a differential AC amplifier (A-M Systems model 1700) and visualized using VScope (Wagenaar, 2017). Timing and relative amplitude of spikes were used to confirm that the observed spikes corresponded to S-cell action potentials while the order of arrival of spikes in the two electrodes was used to determine direction of propagation of action potentials along the nerve cord.

### Mechanical wave generation

A function generator (Pasco Scientific, Roseville, CA) powered two loudspeakers to move a hollow aluminum bar up and down to produce approximately sinusoidal waves in a glass aquarium filled to a depth of 2.4 cm with cold saline. Wave amplitude and frequency were controlled via the function generator. Leeches, immobilized on a PDMS platform and prepared as described above, were placed at a distance of 18 cm from the wave source, in a location where waves propagated approximately linearly. To reduce the influence of visual cues, all experiments were performed in the dark while the room light was turned on between trials to prevent dark adaptation of the animal’s visual system.

To be able to mimic the characteristics of the physical waves in the visual modality (see below), we measured the wavelength of generated waves as a function of frequency, by analyzing the undulations of reflections of brightly lit metal posts placed behind the aquarium. We found that the data were poorly fit by the textbook equation for surface waves, so we used a polynomial fit through the data to generate an experimental dispersion relation. The results of this exercise, for the frequencies used in this study, are displayed in Table 1.

**Table 1.**
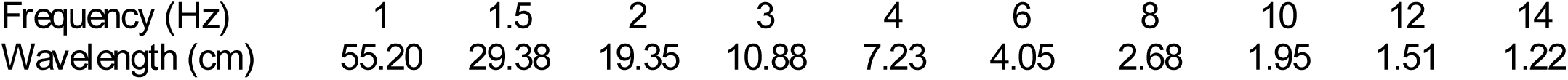
Experimental dispersion relation for mechanical waves

|  |  |  |  |  |  |  |  |  |  |  |
| --- | --- | --- | --- | --- | --- | --- | --- | --- | --- | --- |
| Frequency (Hz) | 1 | 1.5 | 2 | 3 | 4 | 6 | 8 | 10 | 12 | 14 |
| Wavelength (cm) | 55.20 | 29.38 | 19.35 | 10.88 | 7.23 | 4.05 | 2.68 | 1.95 | 1.51 | 1.22 |

### Determination of wave amplitude

To accurately determine wave amplitude for the mechanical experiments, we proceeded as follows. A laser beam was aimed perpendicular to the wave fronts at a downward angle of 45^⁏^. Sawdust was sprinkled onto the surface of the water to illuminate the point where the laser hit the water surface. (We presumed that the weight of sawdust at most marginally impacted the characteristics of the waves being produced, and thus did not influence measurements, but we did not quantitatively verify this assumption.)

Videos of waves were then recorded from an approximately horizontal angle to the water surface. In these videos, brightly lit sawdust particles effectively represented the surface of the water under the laser spot, moving vertically and horizontally with incoming waves. Using custom software (written in Octave) to track bright spots in each video, the movement of the water was measured in the direction of wave propagation and the vertical direction. For each relevant frequency, we measured the wave amplitude at various driving voltages of the function generator to obtain a calibration curve. These curves then allowed us to produce waves of precisely determined amplitudes at all frequencies. At frequencies below 1 Hz, we found that significant bulk water displacement accompanied the surface waves, complicating interpretation of biological responses. Accordingly, we did not include experiments at frequencies below 1 Hz in this paper.

Since it was not possible to image with the camera perfectly horizontal due to the meniscus of the water against the aquarium wall, water movement perpendicular to the direction of wave propagation (towards or away from the camera) could affect the motion estimate in the vertical direction. To quantify this effect, additional videos were recorded with the camera mounted above the aquarium and facing downward onto the water surface. Perpendicular water movements were found to be so small that their effect on vertical measurements was at most 4 µm and hence negligible.

### Visual wave and flash generation

Visual waves were projected from below onto the dorsal surface of a leech using the green channel of a Pico Projector (AAXA Technologies Inc., Irvine, California) and a mirror. All visual waves were square waves with a 50:50 duty cycle of light and dark phases, and were generated in accordance with the experimental dispersion relation of Table 1. Illuminance was controlled using absorptive neutral density filters (Thorlabs, Newton, New Jersey). Experiments were performed under controlled background illuminance of 3.3 ± 0.1 lux of green light.

## Results

Wave stimuli readily evoked spikes in the S cell. This was true for both mechanical waves presented in darkness (Fig. 1A) and for visually cued virtual waves (Fig. 1B). Both modalities elicited a strong initial response followed by a weaker sustained response that was most prominent for low-frequency visual waves (Fig. 1C, D). Cessation of mechanical stimulation but not of visual stimulation occasionally elicited a weak off-response (Fig. 1C). This may have been due to a slight mechanical jerk as the speaker stopped moving.

**Figure 1.**
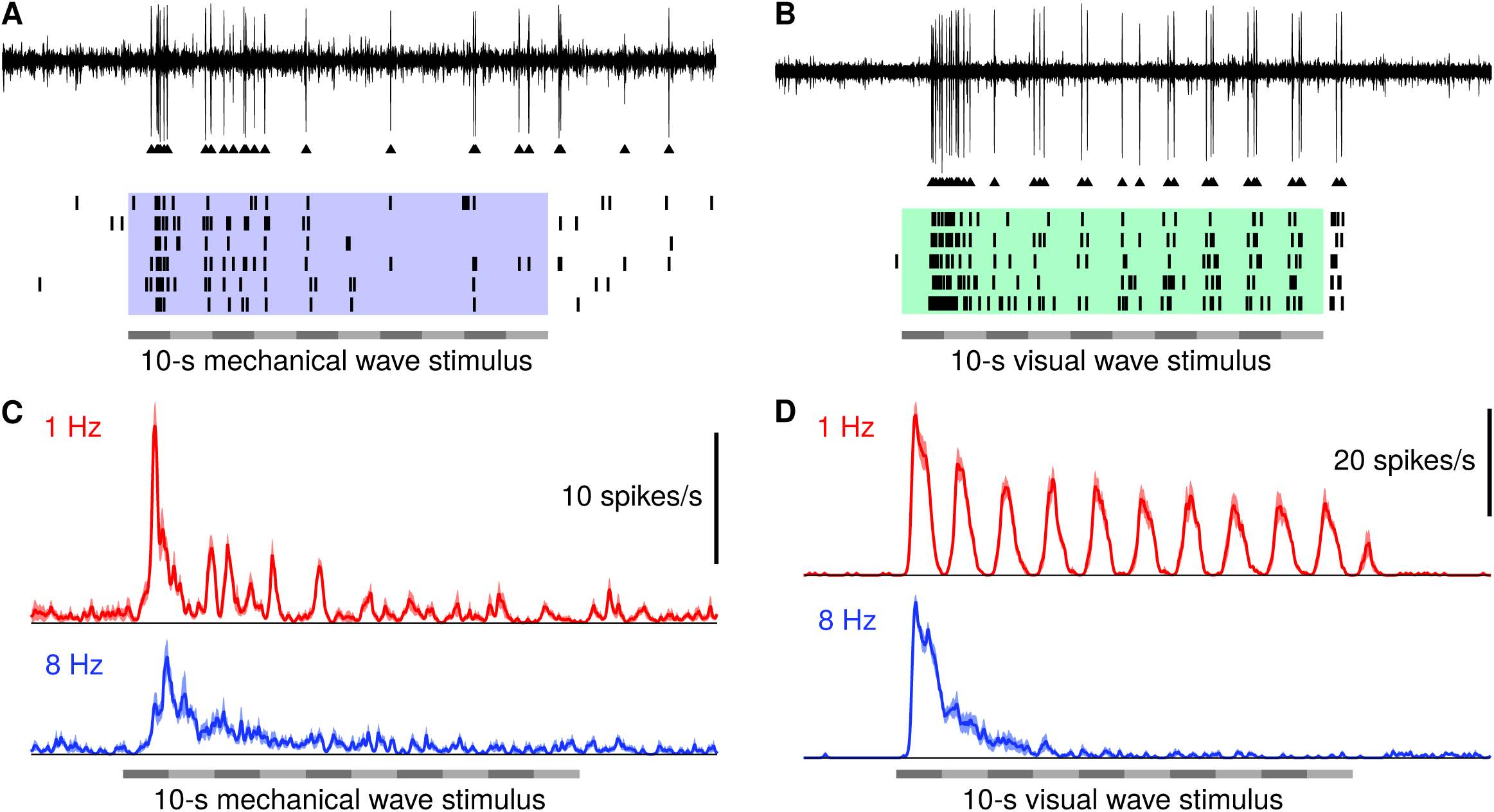
S-cell responses to mechanical and visual wave stimuli. **A.** Extracellular recording (*top*), detected spikes, and raster plot (*bottom*) from an S cell responding to mechanically cued wave stimuli with a frequency of 1 Hz. **B.** Same for visually cued 1-Hz waves. **C.** Average firing rate of the S cell in response to mechanically cued waves at 1 and 8 Hz (mean ± SEM, *N* = 9 leeches.) D. Same for visually cued waves at 1 and 6 Hz (*N* = 6).

Previously, it had been shown that leeches most readily localize mechanically cued wave sources at frequencies around 8 to 12 Hz, whereas they most readily localize visually cued wave sources at frequencies around 2 Hz (Harley et al., 2011). To test whether S-cell activity correlates with these behavioral findings, we exposed leeches to mechanically and visually cued waves of 1 to 14 Hz and varying amplitudes. Regardless of the wave frequency, we found that the S-cell response scaled roughly linearly with wave amplitude for mechanical waves (Fig. 2A). Likewise, the S-cell response scaled roughly linearly with log illuminance (or equivalently: log contrast, as the background illuminance was fixed) for visually cued waves (Fig. 2B). We therefore collected the fitted slopes of the response functions for each of the tested frequencies and used those to define the sensitivity of an individual S cell to waves of that frequency.

**Figure 2.**
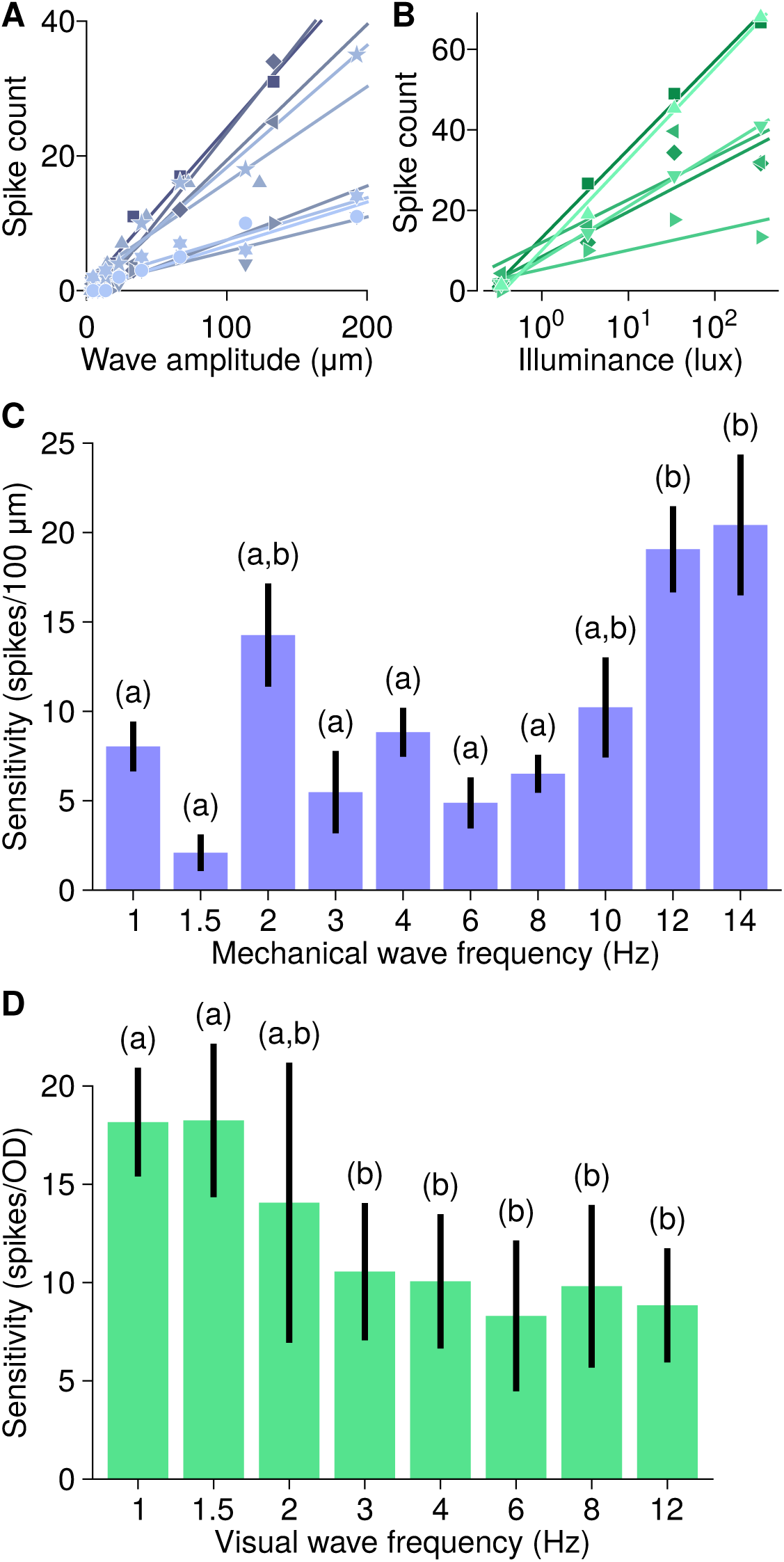
Frequency dependence of S-cell responses. **A.** S-cell spike counts during 10 s of mechanical wave stimulation at 2 Hz, as a function of wave amplitude. Each set of symbols and the corresponding fitted straight line represents data from one of 9 leeches. **B.** S-cell spike counts during 10 s of visual wave stimulation at 2 Hz, as a function of illuminance (*N* = 6). **C.** Sensitivity of the S cell to mechanical waves (see text). Letters indicate groupings from Tukey’s test. **D.** Sensitivity of the S cell to visually cued waves. Letters again indicate groupings from Tukey’s test.

Wave frequency was a significant determinant of the response to mechanical waves (2-way ANOVA with frequency as a fixed factor and animal identity as a random factor; *F*(9, 72) = 9.7; *p <* 10^−**8**^; *N* = 9). Tukey’s test revealed that mechanical waves at 12 and 14 Hz elicited the strongest responses (Fig. 2C), in agreement with behavioral data. For visually cued waves, frequency was also a significant determinant of the response (analogous 2-way ANOVA; *F*(7, 35) = 9.1; *p <* 10^−**5**^, *N* = 6). In contrast to the responses to mechanical waves, responses to visual waves were strongest at low frequencies (Fig. 2D), again in agreement with behavior.

These results suggest that the S cell is involved in prey detection. However, successful prey capture also requires localization of the prey, i.e., determination of the direction of approaching waves. We therefore set out to determine whether the S cell plays a role in this process as well.

We presented leeches with mechanical waves directed toward the head or the tail of the animal or to either the left or right side and measured the S-cell responses. We tested 10 leeches with 1-Hz and 8-Hz waves and counted spikes during the 10 seconds of each stimulus presentation. Although there was a slight trend for lower spike counts in response to waves directed toward the head, ANOVA revealed no significant differences as a function of wave direction at either frequency (Fig. 3A). This was also true when we only counted spikes in either the first 2 seconds of the stimulus or the last 5 seconds (data not shown). We repeated this analysis with visually cued waves, and again found no effect of wave direction (Fig. 3B).

**Figure 3.**
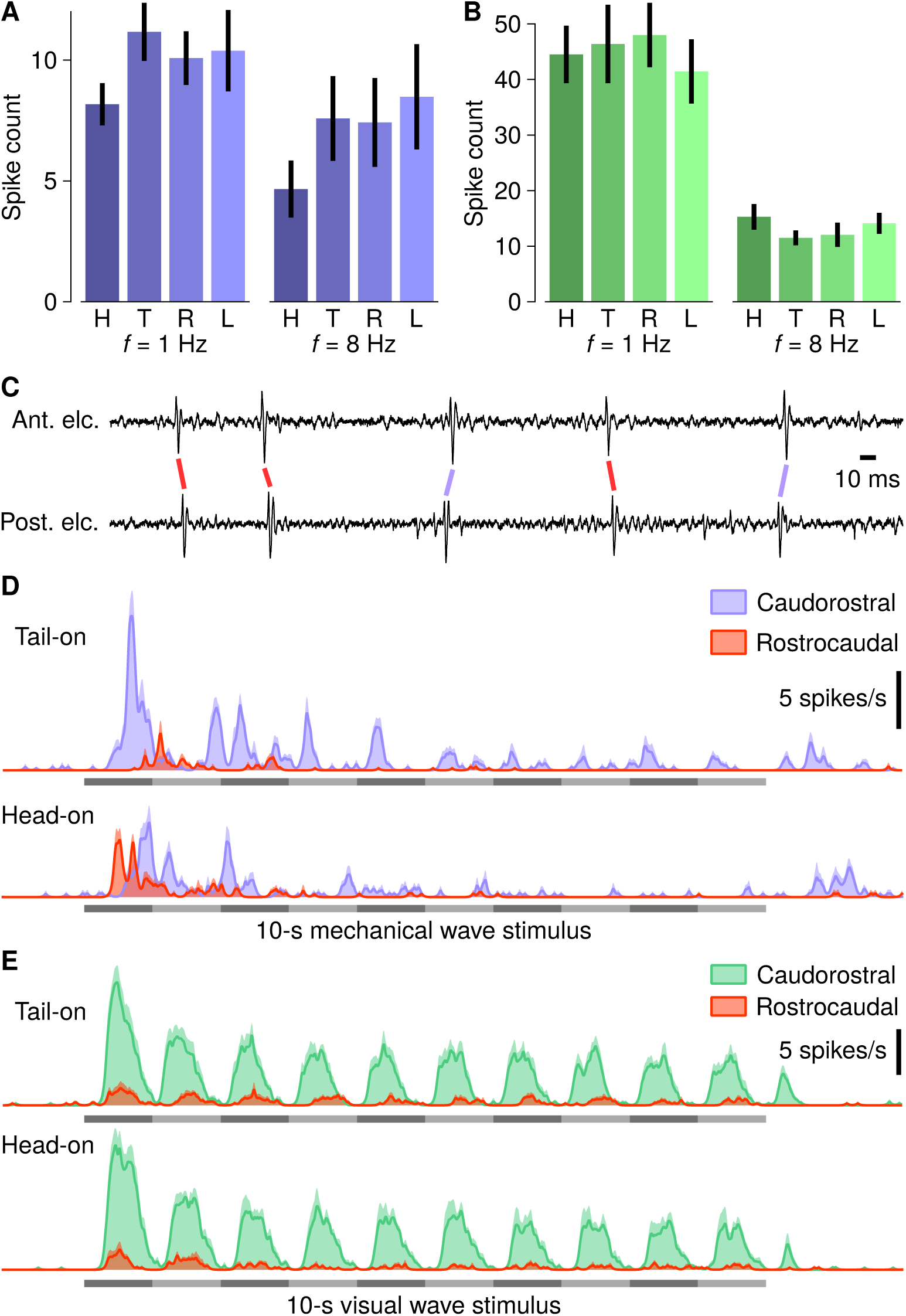
Dependence of S-cell responses on wave direction. **A.** S-cell spike counts during 10 s of mechanical wave stimulation at 1 and 8 Hz for waves directed at the head (H), tail (T), right side (R), or left side (L) of the animal (*N* = 8 leeches). **B.** Same for visually cued waves (*N* = 6 leeches). **C.** Short segment of a recording using two electrodes, one anterior to ganglion 10 (*top trace*), and one posterior to ganglion 10 (*bottom trace*), demonstrating how spike timing can be used to reconstruct propagation direction. **D.** Rate of caudorostrally (tail-to-head; purple) and rostrocaudally (head-to-tail) propagating spikes in response to 1-Hz mechanical waves approaching the animal from the head (*top*) or tail (*bottom*). E. Same for visually cued waves.

However, total spike counts are only part of the story. In all our experiments, we recorded S-cell activity using a pair of electrodes around ganglion 10, which allowed us to not only confirm that the spikes that we recorded were truly S-cell spikes, but also to determine the direction of their propagation (Fig. 3C). This revealed that in response to mechanical waves directed to the tail (“tail-on”), the vast majority of spikes originated posterior to ganglion 10 (Fig. 3D, top), with a small secondary burst of spikes originating on the anterior side shortly after the onset of the posterior response. In contrast, mechanical waves directed to the head (“head-on”), first elicited a strong burst of spikes originating in the anterior, followed by more diffuse spiking from the posterior. In response to waves coming from the left or right side of the leech, the onset latencies of rostrocaudal and caudorostral spiking were equal to each other (data not shown). Remarkably, in S-cell responses to visually cued waves, the direction of wave propagation had no influence on where spikes originated (Fig. 3E). Does this imply that visual information reaches the S-cell system through a more indirect route? To find out, we performed ablation experiments to determine which parts of the nervous system were required for visual responses.

We recorded S-cell responses to full-body light flashes in leeches with intact nervous systems, prior to progressively ablating ganglia posterior to the recording site, to find out whether the caudorostral S-cell spikes originated in the tail brain or in the posterior segmental ganglia. Unlike earlier results, which were all obtained with pairs of electrodes around ganglion 10, here we studied spike propagation around several ganglia (3, 10, 14, or 17). Around ganglia 10 and 14, the vast majority of spikes in response to full-body flashes propagated in a caudorostral direction when the nervous system was intact (Fig. 4A). This was still true when the tail brain (“TB”) and one or two of the most posterior ganglia were ablated, indicating that the tail brain is not required for transmitting visual information to the S-cell system. Sequential ablation of ganglia caused a gradual drop in caudorostral spikes. When almost all of the posterior ganglia were ablated, rostrocaudal spiking increased, indicating that the S-cell system can be activated by visual information in anterior ganglia, but that this doesn’t generally happen when the nervous system is intact, likely because the S-cell system is more readily activated in posterior ganglia. A similar pattern was seen around ganglion 3, although—quite surprisingly—the ablation of even a few posterior ganglia caused an increase in rostrocaudal spiking at this location. Lastly, around ganglion 17, rostrocaudal propagation dominated even when the nervous system was intact—perhaps because there are only four segments posterior to this recording site—and this dominance increased as posterior ganglia were ablated.

**Figure 4.**
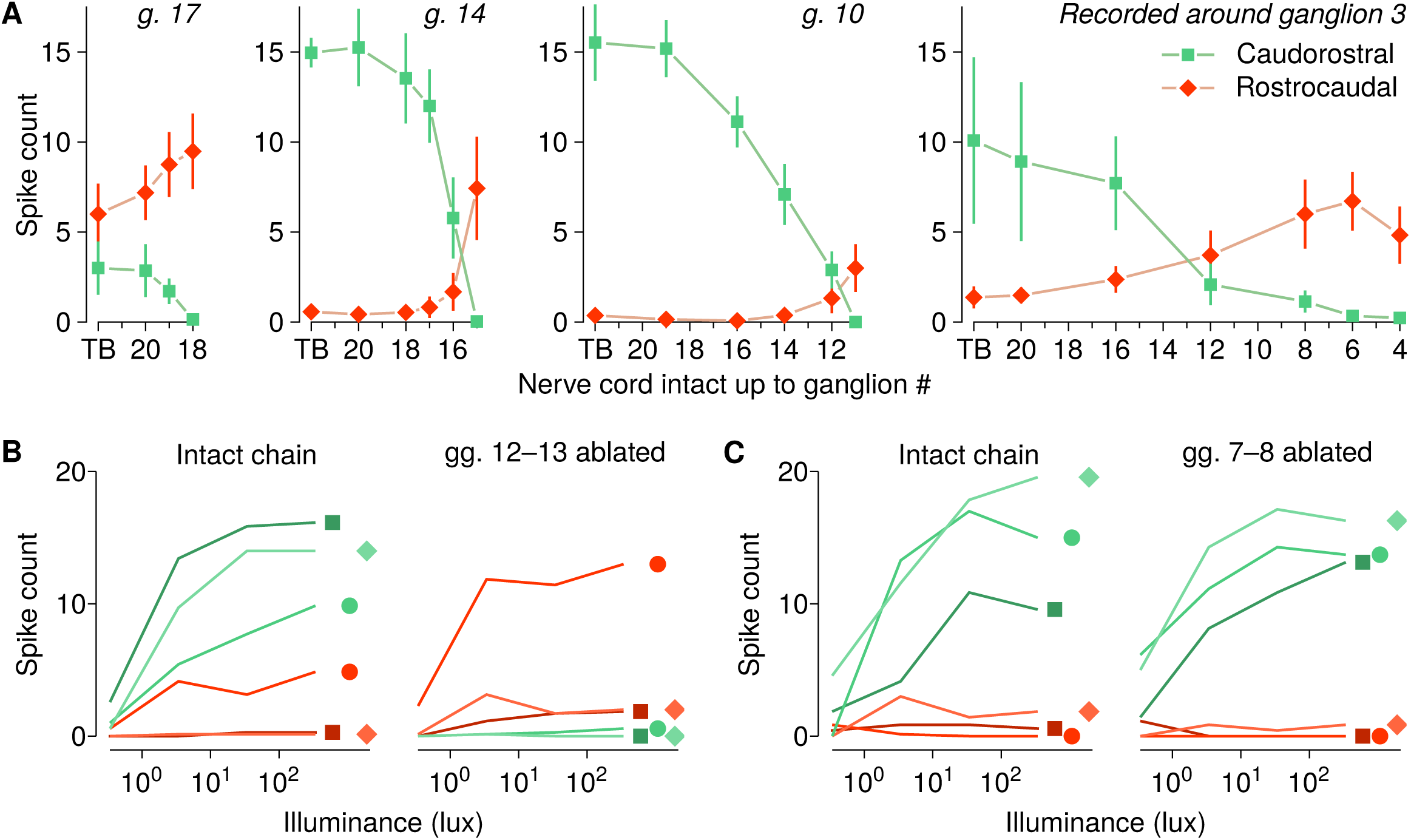
Effects of ablation of portions of the nervous system on S-cell activity. **A.** Counts of caudorostral (*green*) and rostrocaudal (*red*) spikes in response to trains of three flashes (at 1.5 Hz) to the full dorsal surface of the leech at four different sites along the nerve cord in experiments where the posterior portion of the nservous system was progressively ablated. Mean and SEM of spike counts in the 2.5 s following stimulus onset from *N* = 5, 4, 4, 5 leeches at each of the four sites. TB = tail brain. **B.** Counts of caudorostral and rostrocaudal spikes around ganglion 10 in leeches in which the nerve cord was initially transected anterior to ganglion 7 and posterior to ganglion 13 (‘intact chain’), in response to trains of three flashes (at 1.5 Hz) of varying illuminance, and again after ganglia 12 and 13 were ablated. Results from three leeches (*symbols*). **C.** Same as (B), except that ganglia 7 and 8 were ablated.

One more series of experiments was performed to study local propagation of S-cell spikes. Here, we recorded around ganglion 10 as before, but we transected the nerve cord anterior to ganglion 7 and posterior to ganglion 13 to create a chain of 7 ganglia still inside the leech. The nerve roots of ganglia 9–11 were cut as before, so that sensory input could only enter in segments 7–8 and 12–13. We recorded responses to sequences of 3 flashes (at 1.5 Hz) of varying intensity, and found that caudorostral spikes dominated as they did in intact animals (Fig. 4B, left). Ablation of ganglia 12 and 13 caused a dramatic drop in caudorostral spikes (*p <* 0.02, t-test, *N* = 3) and an apparent increase in rostrocaudal spikes in each leech studied, although the t-test did not yield significance. These results indicate that even locally, the posterior segments are a more potent initiator of S-cell spikes. This experiment also demonstrates that the cephalic ganglia are not required for the generation of rostrocaudal S-cell spikes. Ablating ganglia 7 and 8 instead of 12 and 13 had no significant effects on S-cell spiking (Fig. 4C). Likely this was because rostrocaudal spikes were already so rare in baseline that removing their source yielded no noticeable effect.

## Discussion

Hungry leeches are highly motivated to localize the source of water disturbances. Behaviorally, they can use visual cues, mechanical cues, or a combination of the two to infer the presence of water waves and their direction of propagation (Carlton and McVean, 1993; Dickinson and Lent, 1984; Harley et al., 2011, 2013). Little has been known about processing of visual information in the leech, and less about processing of information relating to water movement. It is known, however, that one particular interneuron, the S cell, responds to both visual and mechanical stimuli (Gardner-Medwin et al., 1973; Jellies, 2014; Kretz et al., 1976; Sahley et al., 1994). By presenting nearly intact leeches with natural water waves in complete darkness, we here demonstrated that the S cell responds readily to purely mechanically cued water waves (Fig. 1A, C).

Likewise, by projecting patterns of light and darkness corresponding to propagating waves onto the dorsal surface of the leech, we demonstrated that the S cell responds to purely visually cued water waves (Fig. 1B, D). The responses to low-frequency waves consisted of clear bursts for each passing wave front, whereas at higher frequencies most of the response was in the form of a burst at the onset of the stimulus. What little sustained response there was beyond this initial burst showed little phasic modulation. It is worth noting that in the response to low-frequency mechanically cued waves (Fig. 1C, top) a phase-locked pattern of bursts can also be discerned, but at twice the frequency. Likely, both upward and downward phases of the wave cycle elicited small bursts of action potentials.

The relative sensitivity of the S cell to mechanically cued waves of different frequencies matched the behaviorally measured rate of successful prey localization, and the same was true for the visual modality (Fig. 2). This could either suggest that these response curves reflect properties of the two sensory systems, or that the S cell is directly involved in prey detection. To distinguish between these possibilities, further experiments recording directly from the sensory nerves will be required.

We tested whether the S cell encodes information about the waves’ propagation direction in its firing rate, but found that this was not the case, at any of the wave frequencies tested (Fig. 3A, B). This result makes sense in view of the following two facts: First, there is only one S cell per ganglion, not a pair, and the S cell has a highly symmetric pattern of neurites (Peterson, 1984). This would make differentiation between leftward and rightward propagation difficult. Second, the electrical coupling between S cells in adjacent ganglia is extremely strong (Frank et al., 1975), so that an action potential initiated in any S cell typically propagates through the entire system. This would make differentation between anterior and posterior wave propagation difficult. Nevertheless, we found that the S-cell system does receive directional information about water waves: we found that the direction of spike propagation within the S-cell system depends on the direction of wave propagation, but only when waves are mechanically cued (Fig. 3D, E). Although in all cases the majority of spikes traveled in a caudorostral direction, for mechanically cued waves, the first burst of spikes propagated in the same direction as the waves, suggesting that spikes are initiated in the area where the wave reaches the animal. (The small burst of rostrocaudal spikes after the larger burst of caudorostral spikes in response to water waves directed at the tail could be due to incomplete damping of reflections of the first wave front at the side of the aquarium.) We found that neither the head nor the tail brain are required to generate an S-cell response to visual stimuli. In fact, responses could be elicited in rather short chains of ganglia attached to the local sensory apparatus (Fig. 4B), suggesting that visual information, too, reaches the S-cell system locally in each segmental ganglion rather than through mediation by a remote integration site.

As mentioned, the vast majority of S-cell spikes recorded near the middle of the leech (in segments 10 and 14) traveled in a caudorostral direction (Figs. 3D, E, and 4A). This is likely caused by the same mechanism that causes light flashes onto the tail of a leech to elicit more S-cell spikes than light flashes onto the head (Jellies, 2014; Wagenaar and Stowasser, 2016). It remains unknown whether these effects are due to the posterior sensilla being more sensitive to light and water motion than the anterior sensilla, or due to the coupling between sensory receptors and S cells being stronger in the posterior segments of the leech.

In conclusion, the close match between the tuning curves of the S cell on the one hand, and of the behavioral rate at which leeches can successfully reach the source of a water wave on the other hand, suggests that the S cell plays a role in prey finding. However, since it is unlikely that downstream targets can readout the direction of spike propagation in the S-cell system, the role of the S cells in predation is likely limited to detection of a stimulus and arousal of the animal, and probably does not extend to subsequent localization of the source of that stimulus.

## Acknowledgments

We gratefully acknowledge help from Annette Stowasser in teaching dissection skills and useful discussions on optics and visual systems.

